# First report on Chicken Pediculosis in Modern Battery-Cage Layer Farms in Bangladesh: Behavioral, Pathological and Production Performance Impacts

**DOI:** 10.64898/2026.07.07.737031

**Authors:** Md. Rajiur Rahaman Rabbi, Md. Safowan, Atika Angum Miti, Md. Abdul Mazed Salafi, Md. Zaminur Rahman

## Abstract

The recent shift in Bangladesh from tradition backyard rearing system to modern commercial layer farming has made the birds immune to infectious diseases. Pediculosis, however, continues to pose a challenge in modern production system due to its invasive nature, often going unnoticed and neglected as it is typically non-lethal, yet capable of causing significant production losses. Lice infestation is a persistent threat in poultry production; however, its implications in battery-caged commercial layer hens in Bangladesh remain insufficiently characterized. The study aimed to identify the causative louse species and evaluate its associations with clinical pathology, hematological alteration and productive performance in 30 white-feathered (15 infested + 15 non-infested) and 30 brown-feathered (15 infested + 15 non-infested) laying birds from two commercial farms in Tangail. Morphological characterization confirmed the parasite as *Menacanthus stamineus*, distinguished by a dorsoventrally flattened body, parabolicallly rounded head wider than long, concealed club-shaped antennae, an oblong-oval abdomen with fine setae and three pairs of short legs each bearing paired claws. Infested birds exhibited consistent clinical pathology, including pale combs, petechial hemorrhages around the vent, severe feather damage with alopecic and exudative areas and incidence of irregular and broken-shelled eggs. Production performance analysis revealed significant reduction in hen-day egg production, egg weight, and feed intake, accompanied by significantly increased feed conversion ratios. Hematological evaluation demonstrated significantly reduced hemoglobin concentration, hematocrit and erythrocyte counts in infested hens, indicating mild anemia and compromised oxygen-carrying capacity. Collectively, pediculosis was strongly associated with lice-induced self-inflicted injury and cannibalism, systemic physiological stress, impaired erythropoiesis, reduced production efficiency and compromised welfare in caged laying hens. To best of our knowledge, it was the first integrative reports from Bangladesh documenting *M. stramineus* infestation in battery-caged commercial layer system with concurrent evidence of hematological disruption and measurable productivity losses, underscoring its epidemiological and economic significance and urgent need for targeted, evidence-based ectoparasite control strategies.

## Introduction

Commercial poultry farming in Bangladesh has emerged as a dynamic and rapidly expanding sector of the national economy, contributing in food security, employment generation and rural development. Driven by increasing demand for affordable and safe animal protein, the industry has evolved from traditional backyard rearing systems to modern commercial enterprises equipped with improved housing, biosecurity measures, vaccination programs and advance management practices. About 6 to 8 millions of people directly and indirectly depend on the poultry sector for their livelihoods, including farmers, feed manufacturers, hatchery operators, veterinarians, traders and transport workers (Rahman et al. 2021). With 2.8% annual growth, it contributed to 1.85% and 32.25% of country’s GDP and annual meat supply respectively (Akhter et al. 2009; Kawsar et al. 2013; BBS 2023). The robustness of this sector has been challenged with several constrains such as diseases mainly Avian influenza (AI) and New-castle disease (ND) outbreaks. By contrast, parasitism often neglected as they are rarely lethal in commercial poultry farming due to the recent upgradation in structural and operational biosecurity (Dinka et al. 2010). However, poultry are not free from ectoparasitic infestation due to the invasive nature of the arthropods. External parasites, primarily ticks, fleas, mites and lice, are responsible for growth retardation, lower production performance and life expectancy. A global 44-year (1980–2024) long meta-analysis revealed that the overall prevalence of ectoparasites in chickens was found to be 70.2% (Nahal et al. 2026). Tropical climatic condition in Bangladesh with lack of proper parasite control measures synergize the propagation of those ectoparasites (Naim et al. 2023). Development of insecticidal or acaricidal resistance is a recent addition to this burden (Pu et al. 2024).

Chicken pediculosis, an obligate ectoparasitic infestation, caused primarily by the chewing group of lice such as *Goniodes gigas*, *Menacanthus stramineus*, *Menopon gallinae* and *Lipeurus caponis*, represents a significant health and economic concern in poultry production. Heavy lice infestation adversely affects bird welfare by inducing persistent irritation, preening, feather damage, restlessness, vent picking, anemia and reduced feed efficiency, ultimately leading to poor growth performance and egg production (DeVaney 1976). In severe infestation, birds may experience marked debilitation, immunosuppression and often mortality. Additionally, ruffled feather accompanied by hemorrhage, ulceration and dermal laceration (Materu et al. 2022; Alrahal et al. 2024). The cutaneous damage predisposes the infected birds to secondary opportunistic infections. Furthermore, these ectoparasites may function as vector of various pathogens, thereby exerting detrimental effects on poultry health and overall economic productivity (Mohammad 2020; Saied et al. 2023). In a recent study, the prevalence of lice infestations was 74.2%, and identified species were *M. gallinae* (72.6%), *G. gigas* (11.6%) and *L. caponis* (10.3%) in semi-scavenging indigenous chickens (*Gallus gallus domesticus*) in Bangladesh (Kausar et al. 2025). In addition, infestation with *L. caponis* was also observed in the indigenous chickens of West-Coast India (Narnaware et al. 2025). On the other hand, *M. gallinae* was found in commercial poultry farming in the Enugu state of Nigeria (Mbah-Omeje 2024). Even low levels of chicken body louse (*M. stramineus*) infestations can affect chicken welfare and in a cage-free commercial farming (Murillo et al. 2024). Despite the rapid modernization of poultry industry in Bangladesh, ectoparasitic infestations in intensive commercial layer farming remain largely neglected. Previous studies on chicken pediculosis in Bangladesh mainly focused on backyard or semi-scavenging indigenous chickens, with limited emphasis on commercial cage-based production system (Kausar et al. 2025). Moreover, earlier investigations primarily reported prevalence and species identification, while the behavioral changes, pathological effects and production losses associated with lice infestation have not been adequately explored. To date, no study has yet investigated chicken pediculosis in modern battery-cage commercial layer farms in Bangladesh. Therefore, the study was designed to investigate chicken pediculosis in commercial cage layer farming with emphasis on occurrence, associated behavioral abnormalities, pathological lesions and impact on production performance.

## Methods

### Study design

The study was conducted in two commercial poultry farms located in Sakhipur Upazila, Tangail, Bangladesh and farms were selected based on accessibility, flock size and the willingness of farm owners. The study population comprised layer chickens (NOVOgen^®^) of similar age and production stage. Within each farm, birds were clinically examined and classified into two groups: pediculosis-infested and non-infested. Pediculosis was diagnosed through physical examination of the birds, with careful inspection of common predilection sites including the vent region, under the wings and the neck and dorsal body surface. Approximately 15 birds were selected from each group per farm. Non-infested birds were chosen from the same farm and matched with infested birds based on age, breed or strain, feeding regime and housing conditions. The samples were processed and examined at the Pathology and Parasitology laboratory of the Department of Para-clinical courses, Gono Bishwabidyalay, Savar, Dhaka, Bangladesh.

### Morphological identification of lice

Lice were collected from the infested birds with ethanol (70%) for each host and washed with phosphate buffer saline (PBS). Then the specimens were treated with 10% potassium hydroxide (KOH) to enhance the clarity. After that, the lice were dehydrated and advanced for the preparation of permanent slide. Therefore, parasites were identified using microscope according to the morphological features described by Soulsby (1982) and Wall and Shearer (1997).

### Assessment of production performance

Production performance included hen-day egg production (%), average egg weight (g), daily feed intake (g/bird/day) and feed conversion ratio (FCR) were assessed for both infested and non-infested group birds of each farm.

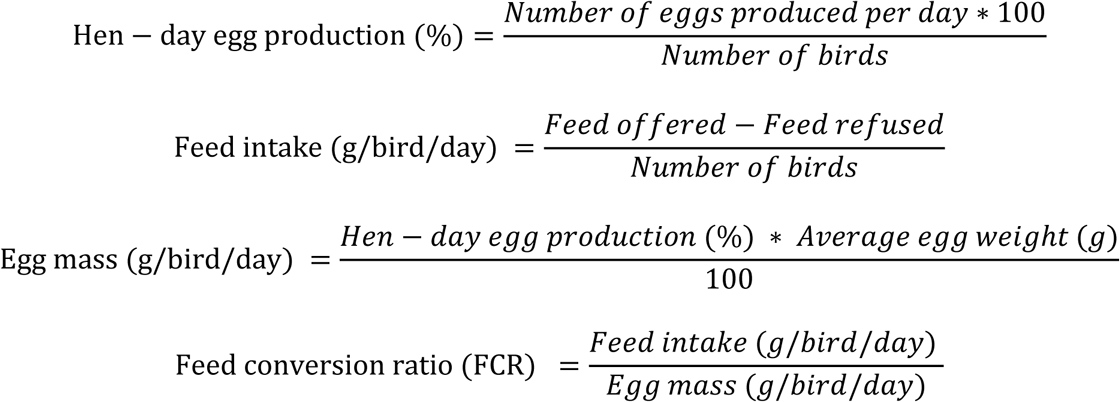

### Haematological examination

Blood samples were collected from the brachial vein of both infested and non-infested birds and then haematological analysis was done using an automatic blood analyzer, BC2800 Vet Auto Haematology Analyzer (Fresenius Medical Care Private Limited, New Delhi). Parameters included Hemoglobin (g/dl), Hematocrit (%) and erythrocyte count (RBCs) (Natt and Herrick 1952; Gaertner and Pazdro 1969; Lamberg and Rothstein’s 1977). Approximately, 1.5 ml of blood sample collected and stored in EDTA tubes for haematological studies.

### Patho-clinical and behavioral observation

The dermis of comb, wattles, vent region, beneath the wings and the neck of the infested group birds were carefully observed for any pathological alteration. For the behavioral observations of the infested birds, particular attention was given to behavioral alterations commonly associated with lice infestations, including restlessness, preening frequency, feather and egg pecking.

### Statistical Analysis

All collected data were entered, managed and analyzed using Microsoft Excel and SPSS (SPSS v25, IBM Corp.). Correlation and independent samples *t*-test were calculated at an 95% confidence level.

## Results

### Demographic data and bird husbandry

Two commercial layer farms located in Sakhipur, Tangail, Bangladesh, were included in the study. Farm 1 housed 50,000 layers(NOVOgen^®^) in six (6) sheds, whereas Farm 2 housed 42,000 layers(NOVOgen^®^) in seven (7) sheds. The selected birds were 74 and 76 weeks of age in Farms 1 and 2, respectively. Cage density was maintained at 3 birds per 2.25 ft² in both farms and sheds were cleaned once daily. No insecticide spray was used in either farm. The selected birds had white feathers in Farm 1 and brown feathers in Farm 2. Both farms followed similar vaccination schedules against Marek’s disease (MD), Herpesvirus of Turkey (HVT), Newcastle disease (ND), infectious bronchitis (IB), infectious bursal disease (IBD), avian influenza (AI), fowl pox, infectious coryza (IC), fowl cholera and egg drop syndrome (EDS). The most recent vaccination administered was against fowl cholera, conducted two weeks prior to sampling in Farm 1 and four weeks’ prior in Farm 2. Feed composition of both farms was consisted of crude protein (18 ± 1%), crude fat (4 ± 1%), nitrogen-free extract (47 ± 1%), crude fiber (≤5 ± 1%), ash (≤15 ± 1%), calcium (≥3.50%), phosphorus (≥0.45%) and metabolizable energy (≥2750 kcal/kg). The maximum feed moisture content was maintained at 12% in both farms.

**Table 1.** Demographic, management and feeding characteristics of the selected commercial layer farms.

|  | Farm 1 | Farm 2 |
| --- | --- | --- |
| **Total no. of birds** | 50,000 | 42,000 |
| **Location of farms** | Sakhipur, Tangail | Sakhipur, Tangail |
| **Bird strain/breed** | NOVOgen<sup>®</sup> | NOVOgen<sup>®</sup> |
| **Total no. of sheds** | 6 | 7 |
| **Time of last vaccination** | 2 weeks ago (Fowl Cholera) | 4 weeks ago (Fowl Cholera) |
| **Cage density(no of birds/sq feet)** | 3/2.25 | 3/2.25 |
| **Cleaning frequency** | Once/day | Once/day |
| **Use of insecticides spray** | No | No |
| **Feather color of selected birds** | White | Brown |
| **Age of selected birds (weeks)** | 74 weeks | 76 weeks |
| **Vaccination** | MD, HVT, ND, IB, IBD, AI, Fowl pox, IC, Fowl cholera, EDS | MD, HVT, ND, IB, IBD, AI, Fowl pox, IC, Fowl cholera, EDS |
| **Feed constituents** |  |  |
| Crude protein (lowest) | 18±1% | 18±1% |
| Crude fat (lowest) | 4±1% | 4±1% |
| Nitrogen free extract (highest) | 47±1% | 47±1% |
| Crude fiber (highest) | 5±1% | 5±1% |
| Ash (highest) | 15±1% | 15±1% |
| Calcium (lowest) | 3.50% | 3.50% |
| Phosphorus (lowest) | 0.45% | 0.45% |
| Energy (lowest) | 2750 Kcal | 2750 Kcal |
| Humidity (highest) | 12% | 12% |

Here, sq feet= square feet; MD = Marek's Disease, HVT = Herpesvirus of Turkey, ND = Newcastle Disease, IB = Infectious Bronchitis, IBD = Infectious Bursal Disease, AI = Avian Influenza, IC = Infectious Coryza and EDS = Egg Drop Syndrome; Kcal= kilo calorie

### Micromorphology of infested lice

The head of the lice was much wider than long and forehead is parabolically rounded. It had club-shaped antennae, which were mostly concealed beneath the head. The abdomen was oblong-oval, elongated and broadly rounded posteriorly and covered with fine hairs (Fig. 1B) and two large setae (Fig. 1A). They had three pairs of short legs with two claws (Fig. 1A). The abdomen is elongated and the last segment is narrowed.

**Figure 1.**
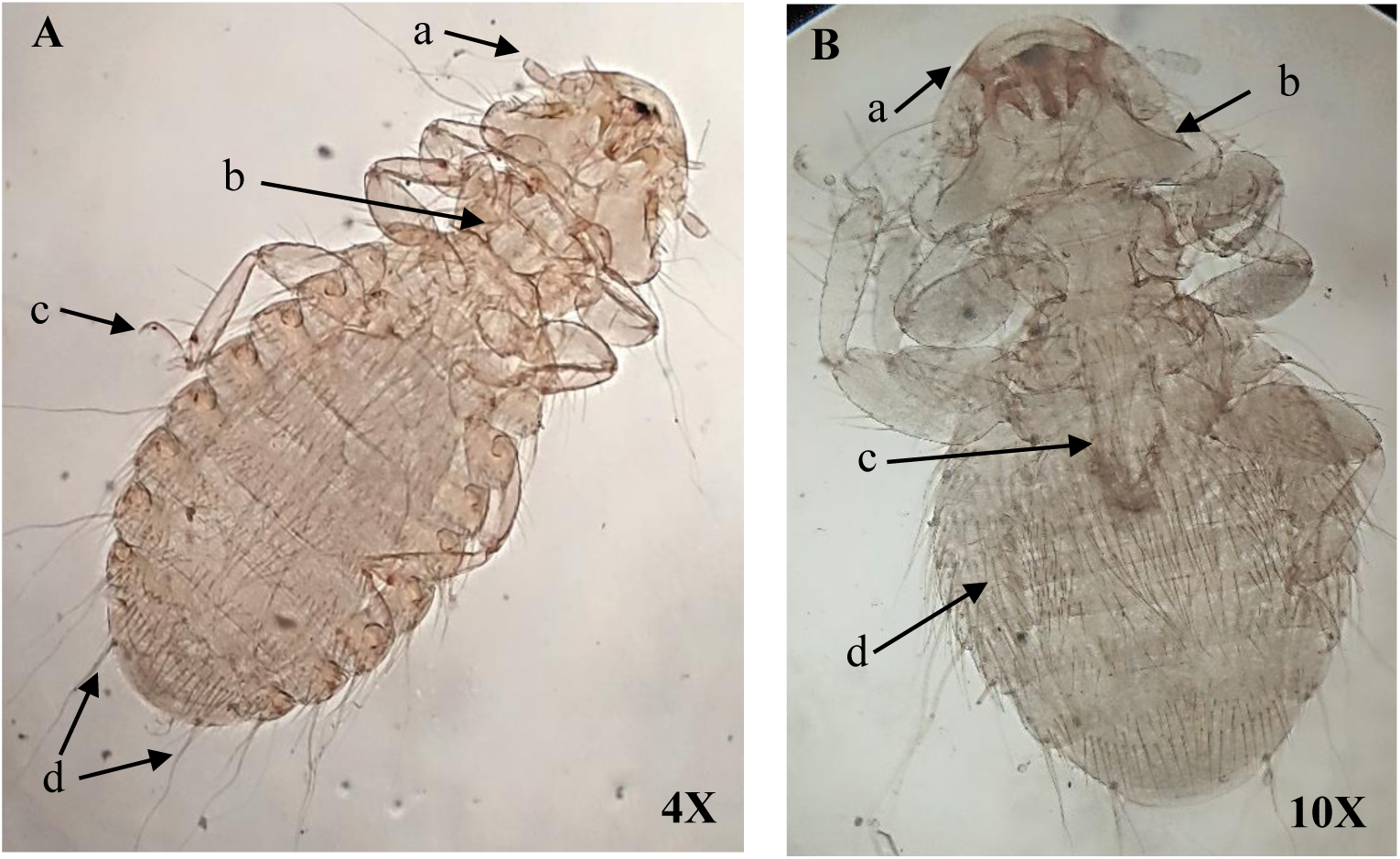
Micromorphology of the body parts of *M. stramineus*. **A.** Adult female *M. stramineus* with club shaped antenna (a), thorax (b), one pair claws in the leg (c) and last abdominal segment containing two large setae at 4X magnification (d). **B.** Nymphal stage of *M. stramineus* with anteriorly rounded head (a), temporal lobe (b), crop (c) and abdominal bristle (d) at 10X magnification.

### Clinical manifestation and pathological changes in infested bird compared to non-infested birds

Compared with non-infested birds, infested birds from both white-feathered (Farm 1) and brown-feathered (Farm 2) birds exhibited noticeably pale combs. Petechial hemorrhagic lesions were observed on the skin surrounding the vent regions, where dense aggregations of brown-colored lice were also detected. Furthermore, infested birds produced eggs with irregular shapes and defective shell quality, including broken-shells. Marked deterioration of feather condition was evident in the affected flocks, characterized by feather loss and alopecic areas with bloody exudation predominantly located on the dorsal and ventral regions.

**Figure 2.**
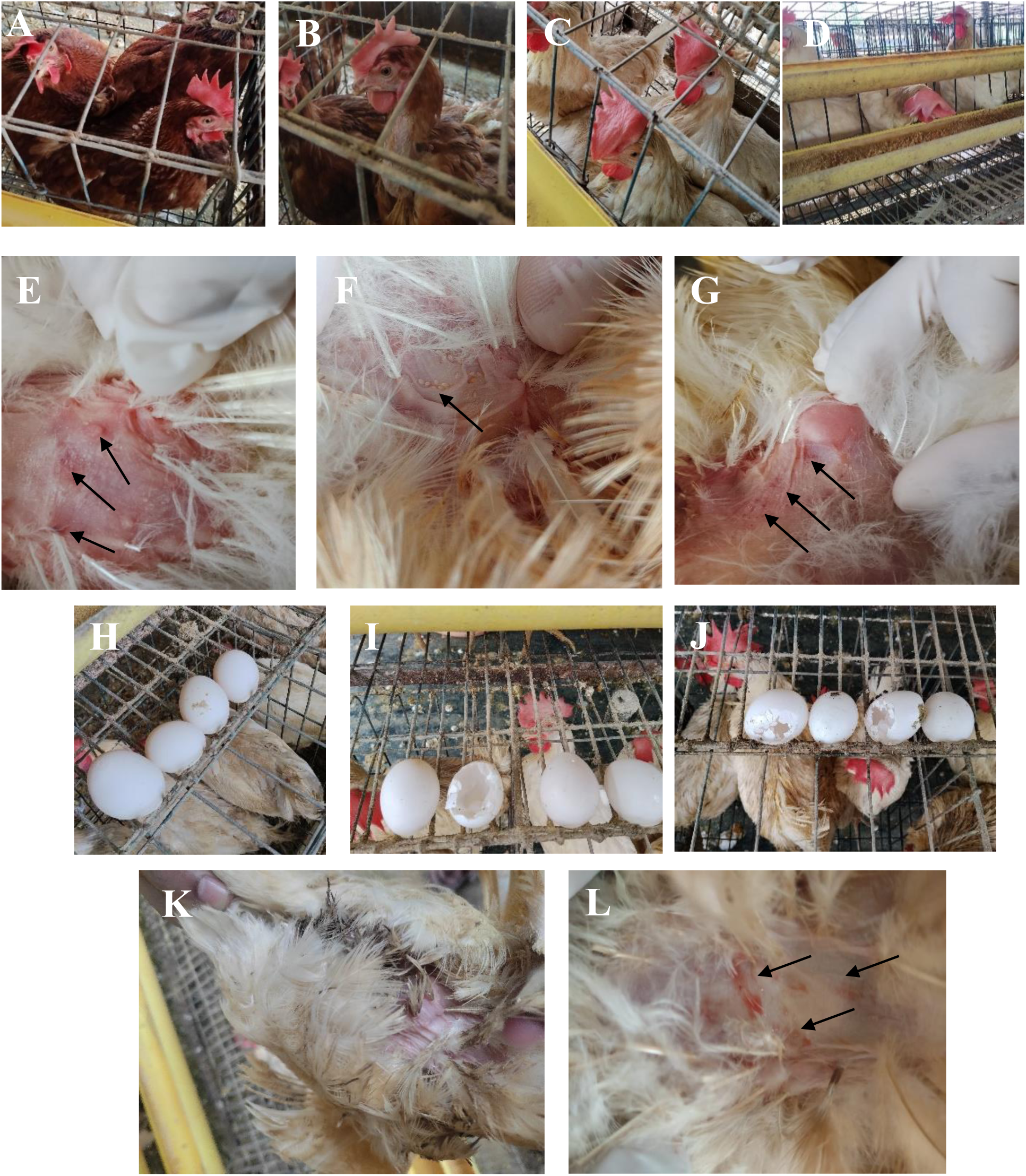
*M. stramineus* induced patho-clinical manifestation in the white- and brown-feathered layers of both selected farms. **A, C.** Non-infested brown-feathered (A) and white-feathered birds (C) with bright reddish color comb. **B, D.** Lice-infested brown-feathered (B) and white-feathered birds (D) with pale color comb. **E, G.** Hemorrhagic foci in the epidermis of vent region. **F.** Colony of *M. stramineus* found near the vent. **H.** Abnormal shaped eggs in infested flocks. **I, J.** Broken eggs in infested flock due to lice-induced peaking by other birds of same cage. **K.** Downgraded feather in infested flock. **L.** Exudation in the vent regions due to lice-induced cannibalism and self-inflicting injury.

### Impact of lice infestation on productive performance parameters

In Farm 1, hen-day egg production was significantly higher (p < 0.001) in non-infested birds (86.06 ± 0.22%) than in infested birds (81.78 ± 0.08%). Similarly, egg weight was significantly greater (p < 0.001) in non-infested birds (59.06 ± 0.57 g) compared with infested birds (52.22 ± 0.51 g). Feed intake was also significantly higher (p < 0.001) in non-infested birds (112.0 ± 1.27 g/bird/day) than in infested birds (104.6 ± 0.74 g/bird/day). In contrast, the feed conversion ratio (FCR) was significantly lower (p < 0.001) in non-infested birds (2.21 ± 0.02) than in infested birds (2.46 ± 0.02). A similar pattern was observed in Farm 2. Non-infested birds exhibited significantly higher (p < 0.001) hen-day egg production (87.95 ± 0.09%) than infested birds (81.22 ± 0.16%). Egg weight was significantly greater (p < 0.001) in non-infested birds (62.20 ± 0.26 g) compared with infested birds (54.15 ± 0.28 g). Feed intake was significantly higher (p < 0.001) in non-infested birds (116.8 ± 0.50 g/bird/day) than in infested birds (107.6 ± 0.79 g/bird/day). Conversely, FCR was significantly lower (p < 0.001) in non-infested birds (2.14 ± 0.01) than in infested birds (2.45 ± 0.02).

**Table 2.** Effect of pediculosis on productive performance parameters of laying hens in selected commercial layer farms.

|  | White-feathered birds (F1) |  | Brown-feathered birds (F2) |  |
| --- | --- | --- | --- | --- |
|  | Non-infested | Infested | Non-infested | Infested |
| Hen-day egg production (%) | 86.06±0.22 | 81.78±0.08*** | 87.95±0.09 | 81.22±0.16*** |
| Egg weight (g) | 59.06±0.57 | 52.22±0.51*** | 62.2±0.26 | 54.15±0.28*** |
| Feed intake (g/bird/day) | 112±1.27 | 104.6±0.74*** | 116.8±0.50 | 107.6±0.79*** |
| FCR | 2.21±0.02 | 2.46±0.02*** | 2.14±0.01 | 2.45±0.02*** |

Here, \*\*\* indicates $p < 0.001$ at 95% confidence level. F1- Farm 1, F2- Farm 2 and FCR- Feed conversion ratio

### Hematological parameters of infested and non-infested white and brown laying hens

In white-feathered birds (F1), non-infested hens exhibited higher hemoglobin concentration (7.57 ± 0.16 g/dl), hematocrit value (28.72 ± 0.14%) and erythrocyte count (2.24 ± 0.01 × 10^6^/µl) compared with infested hens, which showed values of 6.18 ± 0.06 g/dl, 25.89 ± 0.20% and 2.14 ± 0.01 × 10^6^/µl, respectively. Similarly, in brown-feathered birds (F2), non-infested hens had higher hemoglobin concentration (7.50 ± 0.09 g/dl), hematocrit value (28.54 ± 0.19%) and erythrocyte count (2.24 ± 0.01 × 10^6^/µl) than infested hens (6.24 ± 0.09 g/dl, 25.39 ± 0.22% and 2.16 ± 0.01 × 10^6^/µl, respectively).

**Figure 3.**
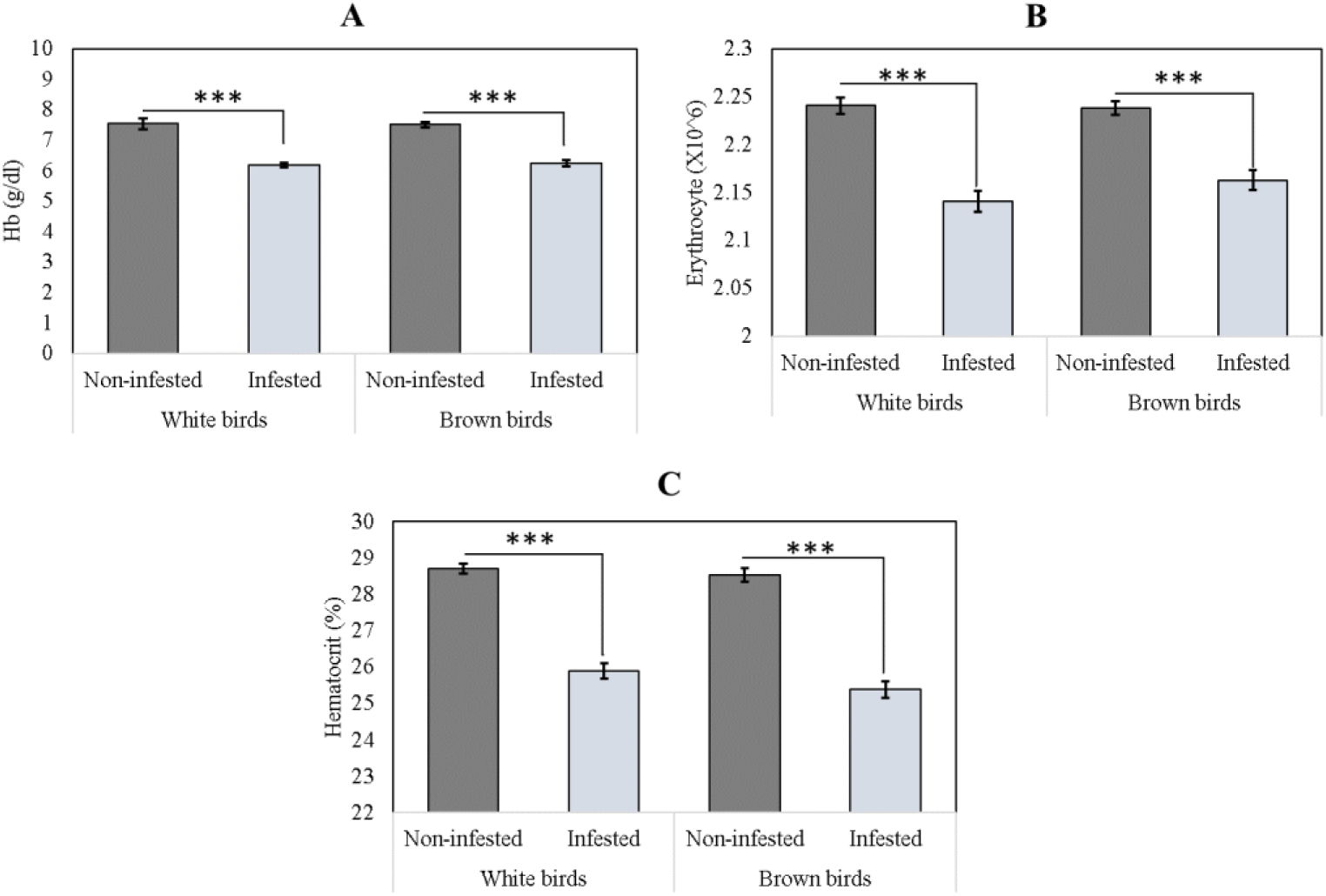
Hematological comparison between infested and non-infested white- and brown-feathered laying hens. **(A)** Hemoglobin concentration (g/dl), **(B)** erythrocyte count (X10^6^µl) and **(C)** hematocrit value (%) in infested and non-infested flocks of selected farms. *** indicates p<0.001 at 95% confidence level.

## Discussion

Lice are among the most prevalent ectoparasite affecting poultry, particularly in backyard and free-range production systems (Endale et al. 2023), although infestations can also occur in intensive farming operations (Mata et al. 2018). Louse infestation can result in substantial economic losses by reducing egg production in laying hens and impairing weight gain in growing birds (Mullens et al. 2010). Infested poultry commonly exhibit severe irritation and pruritis, leading to behavioral changes such as excessive preening, rubbing, restlessness and general discomfort (DeVaney 1976). These particular responses may reduce daily feed intake and negatively affected overall productivity. Furthermore, skin lesions caused by persistent irritation can predispose the layers to secondary bacterial infections. Among the various louse species infesting poultry, the chicken body louse, *Menacanthus stramineus* is considered the most prevalent and economically significant species affecting chickens (Murillo et al., 2014).

The collected lice specimens were identified based on morphological characteristics using the taxonomic keys and descriptions provided by Wall and Shearer (2001) and Yevstafieva (2015). Species identification was performed by examining a range of diagnostic features, including body shape and size, head morphology, antennae position and segmentation, abdominal structure, distribution of body setae (bristles), number of legs and the morphology of the leg claws.

The clinical and pathological finding clearly demonstrated the adverse effects of lice infestation on the health and welfare of laying hens. Infested birds from both farms exhibited pale combs, petechial hemorrhage around the vent region, poor feather condition, alopecic patches, bloody exudation and increased occurrence of damaged eggs compared with non-infested birds. The pale combs observed infested hens are consistent with the reduces hemoglobin concentration, hematocrit values and erythrocyte counts recorded in this study, indicating the development of anemia associated with chronic ectoparasitic infestation. The presence of petechial hemorrhagic lesions and dense aggregations of lice around the vent region suggests continuous irritations and tissue damage caused by parasite activity (Balashov 2007). Persistent irritation likely provoked excessive preening, feather pecking and cannibalistic behavior among cage mates, resulting in the excessive feather loss and alopecic regions with bloody exudate observed in infested birds. Such skin lesion may also predispose affected birds to secondary bacterial infections (Laroche et al. 2018). Furthermore, the higher occurrence of broken-shelled eggs in infested cages may be linked to increased restlessness and cannibalistic or aggressive behaviors induced by lice infestation, leading to accidental egg damage.

The present study demonstrated that lice infestations significantly impaired productive performance in laying hens, as evidenced by reduced hen-day egg production, lower egg weight, decreased feed intake and poorer feed conversion efficiency in both farms. The reduction in egg production and egg weight observed in infested birds may be attributed to the diversion of nutrients and energy from production towards maintenance, immune responses and behavioral activities associated with irritation and discomfort (Murillo et al. 2024). Persistent pruritus and restlessness caused by lice infestation can also reduce feeding time, which likely contributed to the significantly lower feed intake recorded in infested hens (Hinkle and Corrigan 2020). Consequently, inadequate nutrient consumption may have adversely affected egg mass formation and overall productivity. Furthermore, the significantly higher FCR observed in infested birds indicate reduced efficiency in converting feed into egg mass (Li et al. 2024). The finding suggests that lice infestation not only decreases production output but also increases production costs through inefficient feed utilization.

The hematological findings revealed that lice infestation adversely affected erythrocytic profile of laying hens. In both white- and brown-feathered flocks, infested birds exhibited lower hemoglobin concentration, hematocrit value and erythrocyte count compared with non-infested birds. These alterations suggest the development of mild anemia in infested hens, which may be attributed to excessive preening, feather pecking, developed cannibalistic behavior among cage mates, hematophagous nature of lice and the chronic physiological stress associated with infestation (Martin and Mullens, 2012, 2016). Reduces hemoglobin levels and erythrocyte counts indicate a diminished oxygen-carrying capacity of the blood, which can impair metabolic efficiency and overall health status (Cabrales et al. 2006). Likewise, the lower hematocrit values observed in infested birds further support the presence of compromised hematological function. The consistent reduction of these parameters across both farms suggest that lice infestation negatively influences blood physiology irrespective of bird type.

Collectively, the consistent effects observed across both farms highlight the detrimental impact of lice infestation affects bird welfare, layer physiology and flock productivity, emphasizing the need for effective ectoparasite control programs in commercial layer operations. The study acknowledges the limitations of generalizability of the findings as the investigation was conducted in only two commercial layer farms. Therefore, future studies should include larger populations, longitudinal monitoring, molecular epidemiological analyses, economic impact assessments and both in vitro and in vivo evaluations of insecticide efficacy to strengthen evidence-based control strategies for *M. stramineus* in commercial poultry systems. Furthermore, molecular characterization of *M. stramineus* was imperative for the assessment of the role of genetic variation in infestation severity.

## Conclusions

The present study identified *M. stramineus* as the predominant louse species infesting commercial battery-caged layer chickens and demonstrated its significant adverse effects on bird health, welfare, hematological status and productive performance. Infested hens exhibited characteristic clinical manifestations, including pale combs, vent-associated hemorrhagic lesions, feather loss, alopecia, and increased egg damage, accompanied by reduced hemoglobin concentration, hematocrit value and erythrocyte count. Moreover, lice infestation was associated with significant reductions in hen-day egg production, egg weight, feed intake and feed conversion efficiency, highlighting its substantial economic impact on layer production. To the best of our knowledge, this is the first report from Bangladesh documenting *M. stramineus* infestation in battery-caged commercial layer farms together with its associated effects on production performance and hematological parameters. These findings underscore the need for routine surveillance and effective ectoparasite control programs to mitigate production losses, improve bird welfare and enhance the sustainability of commercial poultry production systems.

## Ethics approval

The research was approved by the Animal Ethics Committee, Centre for Multidisciplinary Research, Gono Bishwabidyalay. Savar, Dhaka (CMR/EC/033).

## Acknowledgements

We gratefully acknowledge the Centre for Multidisciplinary Research (CMR) and Department of Para-Clinical Courses, Faculty of Veterinary and Animal Sciences, Gono Bishwabidyalay for its support and the poultry farmers who participated in and facilitated this research.

## Author contribution

Conceptualization, Methods, Data Curation, Formal Analysis, Software, Visualization, Writing (Original draft): Md. Rajiur Rahaman Rabbi. Investigation: Md. Rajiur Rahaman Rabbi, Atika Angum Miti and Safowan Talukder. Writing (reviewing and editing), Resources, Supervision and Validation: Md. Rajiur Rahaman Rabbi and Md. Zaminur Rahman. All authors reviewed and approved the final manuscript.

## Funding

There was no external funding for this particular project, apart from personal contributions by RRR and ZR.

## Data availability

All data generated or analyzed during this study are included in this article.

## Declaration of competing interest

The authors declare no competing interests.

## Notes

### Competing Interest Statement

The authors have declared no competing interest.

